# Predicting Pathogenicity of Missense Variants withWeakly Supervised Regression

**DOI:** 10.1101/545913

**Authors:** Yue Cao, Yuanfei Sun, Mostafa Karimi, Haoran Chen, Oluwaseyi Moronfoye, Yang Shen

## Abstract

Quickly growing genetic variation data of unknown clinical significance demand computational methods that can reliably predict clinical phenotypes and deeply unravel molecular mechanisms. On the platform enabled by CAGI (Critical Assessment of Genome Interpretation), we develop a novel “weakly supervised” regression (WSR) model that not only predicts precise clinical significance (probability of pathogenicity) from inexact training annotations (class of pathogenicity) but also infers underlying molecular mechanisms in a variant-specific fashion. Compared to multi-class logistic regression, a representative multi-class classifier, our kernelized WSR improves the performance for the ENIGMA Challenge set from 0.72 to 0.97 in binary AUC (Area Under the receiver operating characteristic Curve) and from 0.64 to 0.80 in ordinal multi-class AUC. WSR model interpretation and protein structural interpretation reach consensus in corroborating the most probable molecular mechanisms by which some pathogenic BRCA1 variants confer clinical significance, namely metal-binding disruption for C44F and C47Y, protein-binding disruption for M18T, and structure destabilization for S1715N.

## 1 INTRODUCTION

Quickly growing genomic data, largely attributed to next-generation sequencing and high-throughput genotyping, hold great promise for precision medicine. However, a major challenge remains making the stride from genomic variation and other data to diagnostic and therapeutic decision-making. Therefore, there has been a critical need to develop computational methods to predict and understand phenotypic impacts of genetic variants at various biological scales (Cline and Karchin, 2011; Karchin and Nussinov, 2016).

The method development has seen excellent opportunities created by the growing public databases (such as TCGA (Weinstein et al., 2013), dbSNP (Sherry et al., 2001), ClinVar (Landrum et al., 2016) and dbNSFP (Liu et al., 2016)), benchmark studies (Martelotto et al., 2014; Guidugli et al., 2018), and community experiments (in particular, CAGI (Hoskins et al., 2017)). Indeed, some algorithms are widely and successfully applied for functional prediction of genetic variants, such as SIFT (Ng and Henikoff, 2003), PolyPhen2 (Adzhubei et al., 2013), MutationTaster2 (Adzhubei et al., 2013), SNAP (Bromberg and Rost, 2007; Hecht et al., 2015), MutPred (Pejaver et al., 2017a,b), and Evolutionary Action (Katsonis and Lichtarge, 2014, 2017). In addition, the data-driven approach for unraveling genotype-phenotype relationships will continue absorbing the artificial intelligence (AI) and machine learning technologies that have been quickly reshaping other fields (Krizhevsky et al., 2012; Silver et al., 2017).

In this paper, building on our participation in the ENIGMA Challenge in the 5th CAGI experiment, we introduce and assess our novel methods developed for predicting clinical significance and inferring molecular mechanisms for missense variants (single nucleotide variants or SNVs that change resulting amino acids). Specifically, our weakly-supervised machine learning models achieve probability prediction, multiclass classification, confidence (uncertainty) estimation, and mechanistic interpretation of cancer pathogenicity by addressing central questions and making novel contributions in the following two aspects.

First, in the aspect of precision medicine, our central question is how to construct clinical significance predictors that are generally interpretable for diagnosis and potentially actionable for therapeutics. In this study, we combine interpretable and actionable features that describe molecular impacts of SNVs (specifically, impacts on protein structure, dynamics, and function here) and expert-curated labels that accurately summarize clinical significance of SNVs (specifically, the posterior probability of pathogenicity, PoP, and corresponding 5-tier classification by the ENIGMA consortium (Cline et al., to appear)) in supervised machine learning. We examine to what extent molecular-level impacts of SNVs can predict organism- and population-level clinical significance to facilitate the often-expensive clinical phenotyping. We also examine to what extent molecular mechanisms underlying clinically significant SNVs can be uncovered from important features of molecular impacts to help identify potentially actionable therapeutics, which is further probed by structural modeling for some variants.

Compared to some other features commonly used in the field (Martelotto et al., 2014), molecular impacts carry direct causal effects on clinical phenotypes (Pejaver et al., 2017b; Reeb et al., 2016), thus enabling model interpretability; they are available for many genes thanks to relatively inexpensive yet accurate bioinformatics tools (such as MutPred2 (Pejaver et al., 2017b) used in this study), thus enabling broad applicability; and they apply to non-synonymous variants beyond SNVs studied here (such as in-frame or frame-shift indels), thus enabling model generalizability.

Second, in the aspect of machine learning, our central question is how to solve a new type of “weakly supervised” machine learning problems (Zhou, 2018) where the desired label (PoP here) has to be regressed from training data without the exact labels. Such inexact supervision can be observed in many real-world applications where only coarse-grained labels are available because exact labels are too hard or/and too expensive to generate (for instance, in computer vision, annotating bird breeds in images by crowd-sourcing or experts). Particularly in our case, whereas the desired label (PoP) is continuous, the only available labels in the publicly-accessible data are five *ordered* classes which are categorized based on pre-determined PoP ranges (see more details in Sec. 2.1). We develop weakly supervised regressors with tailored loss functions to directly predict PoP. Our first model, a linear one developed during CAGI, used parabola-shaped polynomials for loss functions to penalize predicted PoP values based on their supposed classes (equivalently, PoP ranges here). As the parabola-shaped polynomials as loss functions are too structured and rigid, we continue after CAGI to develop linear and nonlinear (kernelized) models with flexible flat-bottomed loss functions directly learned from data.

Other methods participating in the challenge treat the problem as classification (Cline et al., to appear) and assign PoP afterwards (a non-trivial challenge), among which multi-class logistic regression is compared as a representative in this study. From the perspective of machine learning, they do not consider the order among classes when making classifications (for which a representative of ordinal regression is compared in this study as well) and have to make strong assumptions about the distribution of PoP, albeit implicitly, when converting class category or probability into PoP.

The rest of the paper is organized as following. We first introduce in Materials and Methods the ENIGMA challenge, our training data and feature engineering. We then introduce three types of machine learning models, the first two for multi-class pathogenicity classification whereas the last – weakly supervised regression – newly developed by us for direct prediction of the probability of pathogenicity (PoP). In Results, we start with examining the value of gene type-specific rather than gene-specific data as well as variants data with less confident clinical annotations. We proceed to compare prediction performances among the machine learning models and integrate model interpretation and (protein) structural interpretation to infer molecular mechanisms by which some BRCA1 missense variants could confer pathogenicity.

## 2 MATERIALS AND METHODS

### 2.1 The ENIGMA Challenge

One of the 14 challenges in the 5th CAGI experiment, the ENIGMA Challenge presented 430 BRCA1 and BRCA2 variants (326 exonic and 104 intronic ones) whose clinical significance was newly annotated or recently updated by the ENIGMA Consortium (Cline et al., to appear) and not available in the public domain during the challenge. Specifically, a posterior probability of pathogenicity (PoP) was produced for each variant by multifactorial likelihood analysis (Goldgar et al., 2008) that integrates clinically-calibrated bioinformatics information and clinical information in a Bayesian network. Based on calibrated ranges of PoP shown in Table 1, variants have been classified according to the IARC (International Agency for Research on Cancer) 5-tier classification scheme: Benign, Likely Benign, Uncertain, Likely Pathogenic, and Pathogenic (Classes 1–5). As shown in the Supporting Information (Table S4), this set is skewed to Classes 2 and 1 (Likely Benign and Benign), consistent with the BRCA variants observed in clinical practice.

**TABLE 1.** The 5-tier ENIGMA classification of variants based on the ranges in the probability of pathogenicity (PoP).

| Class | 1. Benign | 2. Likely Benign | 3. Uncertain | 4. Likely Pathogenic | 5. Pathogenic |
| --- | --- | --- | --- | --- | --- |
| PoP range | < 0.001 | 0.001~0.049 | 0.05~0.949 | 0.95~0.99 | > 0.99 |

Participants were asked to predict for each variant PoP according to the ENIGMA classifications as well as confidence level (measured by standard deviation, SD). They were also told that the assessment would be against predicted classes based on PoP ranges (Table 1) instead of PoP and predictions for classes 1 and 5 would be weighted more without the exact formula given. We only submitted predictions for all the 318 missense variants.

### 2.2 Data

To approach the task with supervised learning, we collected missense variants data similarly classified using the five-tier clinical significance system and publicly available in the ClinVar database (Landrum et al., 2016). In other words, no exact values but ranges of PoP (equivalently, classes) are publicly available, creating a weakly supervised learning scenario. The five terms of clinical significance used in ClinVar, following the guidelines from ACMG/AMP (the American College of Medical Genetics and Genomics and the Association for Molecular Pathology) (Richards et al., 2015), are consistent with those used by ENIGMA, following the IARC guidelines. Some ClinVar entries are even submitted by ENIGMA. A slight inconsistency was disregarded, namely that the uncertain range in the ACMG/AMP guideline is 0.10~0.899 rather than 0.05~0.949.

Consistent with the practice during the challenge, during our post-CAGI replication we retrieved missense variants from ClinVar with the last-interpreted date no later than June 29, 2017, around 6 months before the challenge was released. We also set two cutoffs on the review status which ranges from zero to four stars suggesting increasingly reliable interpretation: at least three stars (reviewed by expert panel) or at least two stars (reviewed by multiple qualified submitters without conflict), thus generating two data sets denoted **G2** and **G3** (a subset of **G2**), respectively. The exact query filters used for ClinVar can be found in Table S1 of the Supporting Information (**SI**).

Besides BRCA1/2 variants, we also collected missense variants of other tumor suppressor genes from ClinVar following the same procedure as described above. Even though the challenge was exclusive to BRCA genes, our rationale is to develop predictors that are not just gene-specific but gene type-specific. One benefit from the machine learning perspective is to access more training data and allow for more complex models for accuracy. We used the STRING database(Szklarczyk et al., 2017) to identify other genes whose protein products interact with BRCA1 and BRCA2 proteins and the OncoKB(Chakravarty et al., 2017) database to filter the resulting genes for tumor suppressor genes only. We ended up with 21 other tumor suppressor genes, 17 of which have variant interpretation in ClinVar and are referred to as non-BRCA in short. Details about data-collection procedures and resulting data statistics can be found in the Supporting Information (**SI**) Sec. 1 and 2, respectively.

A stratified split considering the frequencies of the five classes was used to create a held-out ClinVar BRCA test set (one sixth) and a training set (five sixths) from **G2**. The same test set was used when testing **G3**-trained models, whereas the **G3** training set is the subset of the **G2** training set with three stars or more in review status. When other genes are considered, their variants would be added to corresponding training sets.

### 2.3 Feature Engineering

When calculating features for each variant collected for training or validation, we restricted the choices to molecular impacts for interpretable and actionable models. These impacts are often predicted from sequence-level features, which is a great challenge itself. We used MutPred2 (Pejaver et al., 2017b) that predicts the posterior probabilities of loss or gain, whichever is greater, for a wide range of properties induced by amino-acid substitutions. These properties, capturing mutational impacts on protein structure, dynamics, and function, are grouped hierarchically into a custom oncology based on their inherent relationships (Pejaver et al., 2017b). To help model interpretability, we only chose those numeric properties on the 3rd level of the ontology because their parent or child properties are strongly correlated with them whereas themselves are not directly related in calculation and more or less “orthogonal” in molecular mechanism. Therefore, we used as features the posterior probabilities of alteration in 9 properties summarized in Table 2.

**TABLE 2.** The list of properties whose MutPred2-predicted alteration probabilities are used as features.

| Feature Index | Property / Molecular Impact |
| --- | --- |
| 1 | Relative solvent accessibility |
| 2 | Allosteric site |
| 3 | Catalytic site |
| 4 | Secondary structure |
| 5 | Stability and conformational flexibility |
| 6 | Special structural signatures |
| 7 | Macromolecular binding |
| 8 | Metal binding |
| 9 | PTM site |

### 2.4 Machine Learning

#### 2.4.1 Mathematical Description

We consider *n* examples ***x***_*i*_ (*i* = 1,…, *n*) each represented by *q* features, i.e., ***x***_*i*_ ∈ ℝ^*q*^ (∀*i*). Although each ***x***_*i*_ has a continuous label [PoP (probability of pathogenicity), or *p*_*i*_ here], the exact label *p*_*i*_ is not available. Rather, a categorized class of *p*_*i*_ is given according to a customized *K*-tier system: *y*_*i*_ = *k* if *b*_*k* −1_ ⩽ *p*_*i*_ < *b*_*k*_ (*k* = 1,…, *K*) where threshold parameters *b*_*k*_ are increasing in *k.* Without loss of generalizability, *b*_0_ = 0 and *b*_*K*_ = 1, echoing the range of *p*_*i*_. In our study, *K* = 5 and *b*_*k*_’s are set according to Table 1. We use *n*_*k*_ and *c*_*k*_ to denote the number and the portion of examples belonging to Class *k,* respectively.

All vectors are column vectors unless stated otherwise and are denoted in bold-faced lower-case italics. Matrices are denoted in bold-faced upper-case italics.

#### 2.4.2 Models Overview

We describe three types of machine learning models. The first two do not predict PoP (*p*_*i*_) directly but classify pathogenicity (*y*_*i*_) instead: multi-class logistic regression (MLR), a representative multi-class classification method used by other participants in the challenge (Cline et al., to appear), disregard the order among the five classes; and cumulative logit model (CLM) (Agresti, 2003), a representative of ordinal regression, treat the classes as ordered albeit on an arbitrary scale. In contrast, the last, including three “weakly supervised” regressors (WSR) developed by us, directly predict PoP (*p*_*i*_) for variant *i* by training models on inexact labels (not PoP but PoP ranges encoded by pathogenicity class *y*_*i*_) while utilizing both the order and the scale among the classes.

#### 2.4.3 Multi-class Logistic Regression (MLR)

MLR (Böhning, 1992), a multi-class classification model, is an extension of binary logistic regression. We consider a linear model 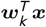 with parameters ***w***_*k*_ for each class *k* (*k* = 1,…, *K*). Let ***w*** denote the column vector stacking all *K* ***w***_*k*_’s. The conditional probability that a sample ***x***_*i*_ belongs to class *k* is thus given by softmax: 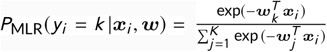. The model parameters 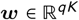 are trained by minimizing the following objective function:

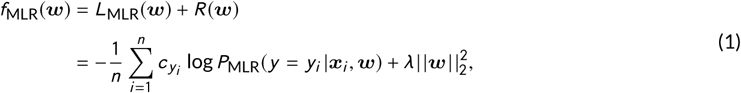

where the loss function *L*(·) is negative weighted log-likelihood, the regularization term *R*(·) is L2 regularization for controlling model complexity, and *λ* is a hyper-parameter for balancing the two terms.

#### 2.4.4 Cumulative Logit Model (CLM)

CLM (Agresti, 2003) is a multi-class classification model that, unlike MLR, considers the order among classes on an arbitrary scale. It is a representative for the type of classification problems called ordinal regression(Gutierrez et al., 2016). Specifically, CLM models the cumulative distribution function by a logistic function *s*(*t*) = 1/(1 + exp(−*t*)): *P*_CLM_(*y*_*i*_ *⩽ k* |***x***_*i*_, ***w***) = *s*(*θ*_*k*_ − ***w***^*T*^ ***x***_*i*_). Therefore, the conditional probability *P*_CLM_(*y*_*i*_ = *k* |***x***_*i*_, ***w***) is now simply the difference between the cumulative distribution functions at two consecutive classes: *P*_CLM_(*y*_*i*_ = *k* |***x***_*i*_, ***w***) = *s*(*θ*_*k*_ − ***w***^*T*^ ***x***_*i*_) − *s*(*θ*_*k* −1_ − ***w***^*T*^ ***x***_*i*_). Compared to *qK* parameters in MLR, a total of *q* + *K* parameters including ***w*** ∈ ℝ^*q*^ and ***θ*** ∈ ℝ^*K*^ are trained here by minimizing a similar objective function (negative weighted log-likelihood with L2 regularization):

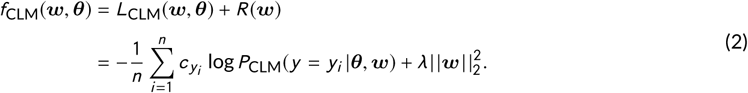

Both baseline models above just calculate the conditional probability that a variant ***x***_*i*_ is classified as each of the *K* = 5 classes: *P*_·_(*y*_*i*_ = *k* |***x***_*i*_, ·). To calculate the probability of pathogenicity *p*_*i*_, we assume that its probability density function is a constant over each class range [*b*_*k* −1_, *b*_*k*_) as specified in Table 1. As such, we estimate *p*_*i*_ by its expectation: 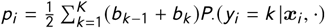.

#### 2.4.5 Weakly Supervised Regression (WSR)

In contrast to using multi-class classification followed by somewhat *ad hoc* assignment of the probability of pathogenicity or PoP (*p*_*i*_ for variant *i*), we developed weakly supervised regression that predicts PoP directly while training data only contains inexact version of PoP, i.e., the pathogenicity class *y* = 1,…, *K.* Specifically, we designed various loss functions 𝓁(*p*_*i*_, *y*_*i*_) that measure the inconsistency between predicted PoP *p*_*i*_ = *p*(***x***_*i*_, ***w***) and its supposed class *y*_*i*_ and trained models by minimizing the L2-regularized loss function:

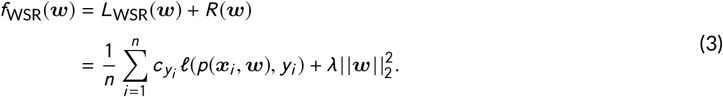

A high-level comparison of the 3 versions of WSR is summarized as following before we describe each in details.

##### WSR1: Fixed, parabola-shaped polynomial loss function

We introduced this model during the challenge and submitted its predictions. Here the PoP predictor is a logistic function *s*(·): *p*(***x***, ***w***) = *s*(*z*(***x***, ***w***)) = *s*(***w***^*T*^ ***x***) whose decision score *z*(·) is linear in ***x***. And the loss function 𝓁(*p*_*i*_, *y*_*i*_) is a pre-determined parabola-shaped polynomial centered around the midpoint of its supposed range 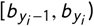:

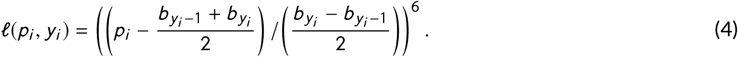

The shape and range in Eq. 3 are very particular, which can be both unnecessary and biased and prevents better accuracy. We thus proceeded to develop much improved models after the challenge, as elaborated next.

##### WSR2: Parameterized *ε*-insensitive loss function

We note that the aforementioned **WSR**1 loss function in Eq. 4 is of a fixed, peculiar shape, which was not optimized during the Challenge and limits predictive power. Therefore, we further developed two more weakly supervised regressors (**WSR**2 and **WSR3**) afterwards. As illustrated in Fig. 1, we allow the class-specific loss function to be flat bottomed (rather than parabola-shaped) in a corresponding parameterized scale (rather than the original, fixed scale) and the label *p* to be transformed (rather than staying in the original space).

**FIGURE 1.**
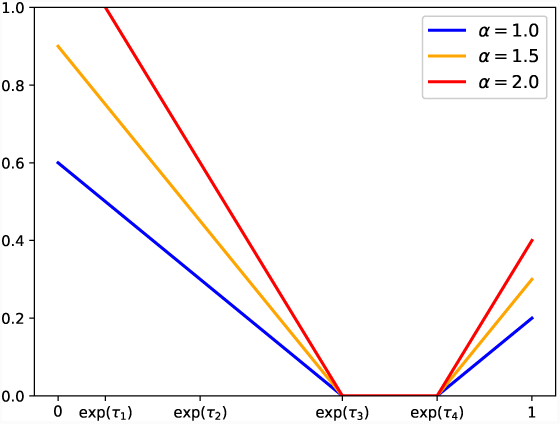
Illustration of the *ε*-insensitive function: max(0, −*α*(*x* − exp(*τ*_3_))) + max(0, −*α* (exp(*τ*_4_) − *x*)) with *α* =1.0,1.5 and 2.0, respectively. The higher the *α* is, the more the penalty the function will give to a prediction outside its supposed range.

Specifically, we first transformed the original scale of fixed thresholds ***b*** over [0, 1] (Table 1) into a new scale of parameterized thresholds exp(***τ*)** (including *τ*_1_ < *τ*_2_ < … < *τ*_*K* −1_ ⩽ 0) to be learned from training data. Note that the use of exp(***τ*)** rather than its logarithm form was designed for the numerical optimization reason (Antal and Csendes (2016)). Correspondingly we had the following transformation between the desired, original label *p*_*i*_ for instance *i* and the transformed label 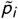 to be predicted:

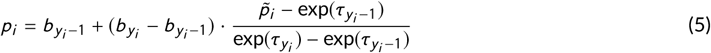

We then used in the transformed 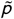-space flat-bottomed *ε*-insensitive loss functions (Vapnik, 2013) that can be regarded as the sum of two hinge loss functions *h*(·). We used class-specific hyperparameters ***α*** to control the slope of a hinge function *h*_*α*_(*x*) = max(0, −*αx*). The higher the *α,* the higher the penalty a prediction would receive if it is outside its supposed range; and no penalty would a prediction receive otherwise. Therefore, for an example ***x***_*i*_ of class *y*_*i*_ with predicted probability 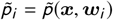, we define its loss as:

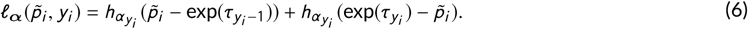

The above expression applies to all classes including the two borders when *y*_*i*_ = 1 or *y*_*i*_ = *K* by introducing constants *τ*_0_ = −∞ and *τ*_*K*_ = 0.

As we did for **WSR**1, we use logistic functions *s*(·) for the predictor of the transformed label 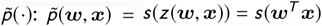 with a linear decision score *z*(·). With the loss for each example redefined in Eq. 6, following the general formula for our weakly supervised regression in Eq. 3, our **WSR**2 models can be learned by solving the following optimization problem:

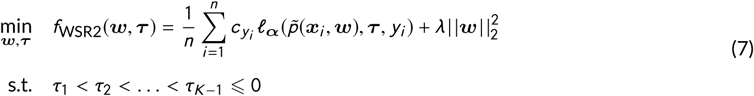

Note that class-specific ***α*** for penalizing out-of-the-range predictions are treated as hyperparameters to be optimized on grid search (see more details in Sec. 2.4.6).

##### WSR3: Kernelized WSR2

The decision score function *z*(·), based on which we used a logistic function to predict transformed label 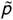, was linear in ***x*** (features) for both **WSR**1 and **WSR**2, i.e., *z*(***w***, ***x***) = ***w***^*T*^ ***x***. Therefore, we further introduce nonlinearity of the decision score function to **WSR2** following a “kernel trick” (Theodoridis and Koutroumbas, 2008) that maps the original feature space to a high-dimensional implicit one and finds linear decision boundary there. Noticing that parameters ***w*** enter the **WSR**2 formulation (Eq. 7) in the form of the inner-product with the feature vector ***x***, but parameters ***τ*** do not, and we have constraints on ***τ.*** We build on the Representer Theorem (Wahba, 1990) (**Lemma 1** in **SI**) and prove a theorem (**Theorem 1** in **SI**) before we apply a kernel trick. A short proof can be found in **SI**.

Based on **Theorem 1**, we can kernerlize the objective function in the **WSR**2 formulation (Eq. 7) to reach the following formulation for **WSR**3

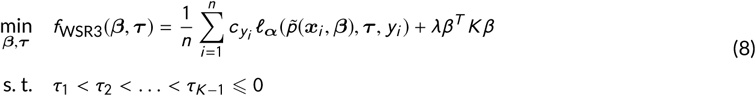

where 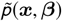 is now logistic in the kernel space, i.e., 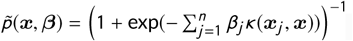.

As such, the kernel trick maps the original feature space ***x*** into an infinite-dimensional space without the need of calculating the exact mapping between them. This trick will enable our model to deal with the non-linear situation, which could significantly increase our model accuracy. In this paper, we use radius basis function (RBF) kernels with bandwidth *γ*. The hyperparameter *γ* is optimized by cross-validation with more details given next.

#### 2.4.6 Model Training and Uncertainty Estimation

In order to obtain the uncertainty measure we randomly split the training set into 5 folds and trained 5 models on 5 combinations of 4 folds. This random split was fixed across all types of machine learning models in the study. The predictions of the five models on the test set are used to calculate the mean and the standard deviation of both the label and the assessment metrics. We then trained on all the 5 folds of the training set to obtain the final model for interpretation.

The hyperparameters are optimized using grid search through 4-fold or 5-fold cross validation depending on the number of folds included in the training set. Specifically, the grid for regularization coefficient *λ* consists of 25 points, for which the log2 of them are uniformly distributed on [−3, 4]. We use this grid for the regularization constant in all models mentioned before. For **WSR**2 and **WSR**3, class-specific slope of the *ε*-insensitive loss function, *α*_*i*_, is sampled on the grid of 4 points: [2^−0.5^, 2^0^, 2^0.5^, 2^1^]. For **WSR**3, the bandwidth of the RBF kernel, *γ*, is sampled on the grid of 25 points, for which the log2 values are evenly distributed between −3 and 2.

For each combination of hyperparameter values, we find model parameters by solving the corresponding optimization problem. Except that **MLR** (multiclass logistic regression) is already implemented in Python-sklearn(Pedregosa et al. (2011)), we implemented all other models in Python 2.7 with optimizers provided in SciPy (Jones et al., 2001–). **CLM** (cumulative logit model) involves a convex optimization problem and was solved by the optimizer ‘BFGS’.

**WSR**1 (weakly supervised regressor 1) involves a nonconvex, unconstrained optimization problem and was solved by BFGS with multi-start. Similarly, **WSR**2 and **WSR**3 involve nonconvex, constrained optimization problems and were solved by the optimizer ‘L-BFGS-B’ with multi-start. Specifically, we sampled 100 initial coordinates for ***w*** and ***τ,*** where *w*_*i*_ s are uniformly sampled from [−5, 5], and *τ*_1_, *τ*_2_, *τ*_3_, *τ*_4_ are uniformly sampled from [−10, 0] with the constraint of *τ*_1_ < *τ*_2_ < *τ*_3_ < *τ*_4_. The rationales for these ranges are the following. First, the standard logistic function *s*(***w***^*T*^ ***x***) saturates quickly when |***w***^*T*^ ***x***| becomes large and, since our features ***x*** are all MutPred2 property probabilities between 0 and 1, large *w*_*i*_ would push transformed PoP values 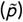 close to 0 or 1. Second, since exp(*τ*_*i*_) is a threshold in the scale of 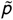, the exponential function exp(*τ*_*k*_) will quickly saturate when *τ*_*k*_ becomes far negative, which pushes all *τ*_*k*_’s to become equal. We analyze the detailed optimization results in **SI** Sec. 4.

#### 2.4.7 Model Interpretation

We further analyze feature importance for each pathogenic or likely pathogenic variants (Class 4&5) in the ENIGMA Challenge (denoted the set of *S*_1_), based on the prediction of model **WSR**2 for its balance of accuracy and interpretability. In other words, for *each* variant *i* ∈ *S*_1_, we would like to examine the statistical significance by which each feature actually separates the variant from the set of benign and likely benign examples (Class 1&2) in the ENIGMA Challenge (denoted the set of *S*_0_). From Eq. 5 and logistic function, we know that the PoP is monotonically increasing with respect to the decision score *z* = ***w***^*T*^ ***x***: the higher the decision score the variant has, the more pathogenic it is. Therefore, given all benign or likely benign variants’ decision scores *z*_*j*_ = ***w***^*T*^ ***x***_*j*_ (1 ⩽ *j* ⩽ |*S*_0_ |), for each variant *i* ∈ *S*_1_ with decision score *z*_*i*_ = ***w***^*T*^ ***x***_*i*_ (∀*i* = 1,…, |*S*_1_ |), we calculate the following differences for the *r*-th feature: 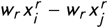 (where *i* is given and *j* ∈ *S*_1_. With the differences defined above, we perform the one-tailed one-sample *t*-test for each variant *i*’s feature *r* and calculate corresponding P-values, where the null hypothesis is that the expected value of the differences, treated as a random variable, is bigger than 0. Finally, for each variant *i* ∈ *S*_1_, We rank all its features based on their P-values in an increasing order, where the smallest P-value corresponds to the most important feature.

### 2.5 Assessment Metrics

Due to the unavailability of ground-truth PoP values, we use two metrics to assess multi-class classification performances. The first is the multi-class area under the curve (AUC) of receiver operating characteristic (ROC) curve (Hand and Till, 2001), which is simply the unweighted average of binary AUC between all pairs among *K* classes:

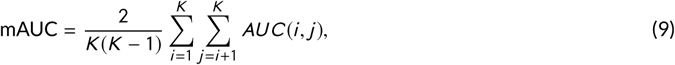

where *AUC*(*I*,*j*) denotes the binary AUC between class *i* and *j.*

As the metric disregards orders among classes, we adopted a second metric, ordinal *mAUC,* which estimates the joint probability that randomly picked instances, one from each class, are scored in the supposed order (Waegeman et al., 2008).

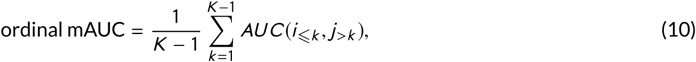

where *AUC*(*i*_⩽*k*_, *j* _>*k*_) denotes the binary AUC between two partitions of all classes: the first *k* and the rest.

As the official assessment did (Cline et al., to appear), we also assess performances on binary classification when classes 1 and 2 are merged to a negative class and classes 4 and 5 are merged to a positive. We use binary AUC as well as RMSD (root mean squared deviation) in PoP. Since ground-truth PoP values are not available, they are approximated to be 0.025 and 0.975 in the assessment.

### 2.6 Protein Modeling for Structural Interpretation

We investigate the impact of pathogenic misssnese mutations on BRCA1/2 proteins through structural modeling. There are only ten pathogenic BRCA variants (class 5) in the ENIGMA Challenge (also referred to as the CAGI test set) including seven for BRCA1 and three for BRCA2. Furthermore, among these pathogenic variants, only five BRCA1 variations occur in available 3D structures (Bateman et al., 2017), including four in the RING (Really Interesting New Gene) domain (M18T, C44F, C47Y, R71G) and one (S1715N) in the BRCT (The BRCA1 C-terminal) domains.

Following various mechanistic hypotheses (such as mutational impacts on folding stability and binding affinity), these five variants are structurally modeled by re-designing wild-type structures using multi-state protein design method iCFN (Karimi and Shen, 2018). Residues within 5 Å from the mutation site were allowed to be flexible (as discrete rotamers) in all designs except that they were extended to those within 8 Å when modeling the mutational effect of S1715N on BRCT-protein binding as the mutation site is at the second layer of the binding interface.

For modeling the effect of four RING-domain mutations (M18T, C44F, C47Y, R71G) on folding stability, we used the single state design (positive only) with substate ensembles where substates were defined as the BRCA1 RING domain in 14 NMR structures in complex with BARD1 (PDB ID: 1JM7). Substate energies were folding energies of the RING domain only that include Coulomb electrostatics, van der Waals, internal energies (Geo term), and a nonpolar contribution to the hydration free energy based on solvent accessible surface area (SASA) (Shen et al., 2015, 2013; Shen, 2013). A positive-substate stability cutoff and positive-versus-negative substate specificity were essentially not mandated with a cutoff of 1,000 kcal/mol.

For modeling the effect of the four RING-domain mutations on interactions with BARD1 (RING domain as well), we used multi-state design (positive and negative) with protein-complex substate ensembles defined in the same PDB entry 1JM7. Positive substate energies were the total folding energies of the RING domain and BARD1 separately and negative substate energies were folding energies of the complex of RING domains of BRCA1 and BARD1. A positive-substate stability cutoff was set at 10 kcal/mol and positive-versus-negative substate specificity was essentially not mandated with a cutoff of 1,000 kcal/mol.

Similarly, for modeling the effect of one BRCT-domain mutation (S1715N) on the stability and protein interaction of BRCT, we did the same as described above except that there was only a single substate available in the crystal structure. For modeling protein interaction, we used BRCT interacting with Bach1 Helicase (PDB ID: 1T29) and for modeling the stability, we used the unbound structure of BRCT domain (PDB ID: 1JNX).

For either folding stability or binding affinity, top conformations of each designed sequence in each substate (backbone conformation here) generated from iCFN for either state were geometrically grouped into representatives. Later, folding stabilities (*G*) and binding affinities (Δ*G*) of the top sequence-conformation ensembles were re-evaluated and re-ordered with a higher-resolution energy model where continuum electrostatics replaced Coulombic electrostatics (Karimi and Shen, 2018). Lastly, the representative conformation at either state was chosen based on the best binding affinity or folding stability (lowest Δ*G* or *G*) for modeling the binding or folding respectively. Each calculated relative binding energy to wild type (WT), ΔΔ*G,* and relative folding energy to WT, Δ*G,* was further decomposed into contributions of van der Waals (vdW), continuum electrostatics (elec), SASA-dependent nonpolar solvation interactions (SASA), and internal energy (Geo).

## 3 RESULTS

### 3.1 The Value of Gene Nonspecific Data and Less Confident Clinical Annotations

We first assess the value of more abundant albeit less confident variant annotations. When restricted to those at least reviewed by panel (‘review status’ being at least 3 stars, or **G3**), the number of BRCA variants accessible from ClinVar was 201 (Table S2 in the Supporting Information), which increased to 699 (Table S3) when the review status was relaxed to at least 2 stars, or **G2** (at least reviewed by multiple qualified submitters without conflict). The larger **G2** training set, even though their annotations are less confident, actually led to no worse multi-class logistic regressor (MLR). As shown in Table 5, mAUC (ordinal mAUC) was improved by 6% (4%) for the ClinVar test set and even more – 8% (9%) – for the CAGI test set in the posterior analysis, respectively.

We also examine to what extent a gene non-specific predictor can rival a gene specific one for pathogenicity prediction. We focus on a gene-type specific predictor here by restricting to 17 other tumor suppressor genes interacting with BRCA1/2. There were 895 **G2** non-BRCA variants much more heavily skewed toward Class 3 (uncertain significance) compared to BRCA variants in the ClinVar set (Table S3). The BRCA variants in the CAGI set turned out to be heavily skewed toward Class 2 (likely benign) instead. Compared to that trained on the BRCA-only data, the MLR model trained on the non-BRCA data had slightly worse performance (8%~11%), whereas that trained on both data (nearly 60% are non-BRCA variants) performed equally.

These results indicate the value of less confident albeit more abundant variant data for pathogenicity prediction. They also show the promise of gene type-specific pathogenicity predictors that can access more variant data of more genes and allow for more complex machine learning models with more parameters. We thus use the **G2** dataset with all the 19 genes thereinafter.

### 3.2 Comparing Machine Learning Models

We next compare three types of machine learning models, as previously summarized in Table 3, for the task of pathogenicity prediction: multi-class logistic regression (MLR), a representative of multi-class classification used by other participants in the Challenge (Cline et al., to appear); cumulative logit model (CLM), a representative of multi-class classification that considers the order among classes (ordinal regression); and our three weakly supervised regression (WSR) models that directly predict the probability of pathogenicity from training pathogenicity classes using designed loss functions, summarized and comapred in Table 4. Due to the lack of ground-truth PoP values, we were only able to assess classification performances. The comparison using multi-classification metrics (mAUC and ordinal mAUC) is given in Table 6 whereas that using binary classification metrics (binary AUC and RMSD) can be found in Table 7.

**TABLE 3.** Overview of two types of baseline machine learning models compared in this study as well as our weakly supervised regression.

| Model | Target Label | Learning Type | Ordered Classes | Scaled Classes |
| --- | --- | --- | --- | --- |
| Multi-class Logistic Regression | Pathogenicity Class | Classification | ✗ | ✗ |
| Cumulative Logit Model | Pathogenicity Class | Classification | ✓ | ✗ |
| Weakly Supervised Regression | Pathogenicity Probability | Regression | ✓ | ✓ |

**TABLE 4.** Comparing the three versions of weakly supervised regression (WSR) models on their loss functions, thresholds, label transformation, and feature embedding.

| Model | Loss function $\ell(p_i, y_i)$ | Threshold | Label Transformation | Feature Embedding |
| --- | --- | --- | --- | --- |
| WSR1 | Fixed polynomial | Constant $\mathbf{b}$ | × | Linear |
| WSR2 | Parameterized ε-insensitive | Parameter $\tau$ | ✓ | Linear |
| WSR3 | Parameterized ε-insensitive | Parameter $\tau$ | ✓ | Nonlinear (Kernelized) |

**TABLE 5.** The classification performance of multi-class logistic regression trained on various datasets.

| Dataset for Training | Clinvar Test Set |  | CAGI Test Set |  |
| --- | --- | --- | --- | --- |
|  | mAUC | Ordinal mAUC | mAUC | Ordinal mAUC |
| **G3** with **BRCA** only | 0.572 ± 0.025 | 0.588 ± 0.017 | 0.532 ± 0.034 | 0.512 ± 0.015 |
| **G2** with **BRCA** only | 0.639 ± 0.009 | 0.623 ± 0.005 | 0.610 ± 0.004 | 0.603 ± 0.011 |
| **G2** with non-**BRCA** genes | 0.542 ± 0.010 | 0.535 ± 0.008 | 0.503 ± 0.014 | 0.521 ± 0.008 |
| **G2** with all genes | 0.640 ± 0.012 | 0.632 ± 0.008 | 0.611 ± 0.010 | 0.636 ± 0.006 |

**TABLE 6.** Pathogenicity-prediction performance comparison (5-class evaluations) among **MLR** (multi-class logistic regression), **CLM** (cumulative logit model), and our **WSR** (weakly supervised regression) variants.

| Model | ClinVar Test Set |  | CAGI Test Set |  |
| --- | --- | --- | --- | --- |
|  | mAUC | Ordinal mAUC | mAUC | Ordinal mAUC |
| **MLR** | 0.640 ± 0.012 | 0.632 ± 0.008 | 0.611 ± 0.010 | 0.636 ± 0.006 |
| **CLM** | 0.673 ± 0.016 | 0.691 ± 0.005 | 0.635 ± 0.009 | 0.684 ± 0.026 |
| **WSR1** | 0.646 ± 0.016 | 0.682 ± 0.012 | 0.623 ± 0.011 | 0.666 ± 0.008 |
| **WSR2** | 0.754 ± 0.015 | 0.782 ± 0.011 | 0.763 ± 0.010 | 0.778 ± 0.023 |
| **WSR3 (Kernelized WSR2)** | 0.791 ± 0.008 | 0.826 ± 0.014 | 0.781 ± 0.010 | 0.802 ± 0.017 |

**TABLE 7.** Pathogenicity-prediction performance comparison (2-class evaluations) among **MLR** (multi-class logistic regression), **CLM** (cumulative logit model), and our **WSR** (weakly supervised regression) variants.

| Model | ClinVar Test Set |  | CAGI Test Set |  |
| --- | --- | --- | --- | --- |
|  | Binary AUC | RMSD | Binary AUC | RMSD |
| **MLR** | 0.751 ± 0.004 | 0.251 ± 0.005 | 0.720 ± 0.011 | 0.277 ± 0.003 |
| **CLM** | 0.853 ± 0.006 | 0.220 ± 0.004 | 0.854 ± 0.008 | 0.213 ± 0.005 |
| **WSR1** | 0.801 ± 0.010 | 0.212 ± 0.012 | 0.780 ± 0.005 | 0.234 ± 0.010 |
| **WSR2** | 0.971 ± 0.001 | 0.201 ± 0.004 | 0.961 ± 0.003 | 0.198 ± 0.004 |
| **WSR3 (Kernelized WSR2)** | 0.982 ± 0.003 | 0.131 ± 0.004 | 0.968 ± 0.002 | 0.161 ± 0.003 |

By comparing **MLR**, **CLM**, and **WSR**1, we found that they had very similar performances for both the ClinVar and the CAGI sets in mAUC that disregards orders among classes; and **CLM** and **WSR**1, both respecting class order in their models, improved for both sets ordinal mAUC that addresses class order. **WSR**1 did not improve against **CLM**, a representative ordinal regression model, because **WSR**1 suffers from its very peculiar loss function that is fixed to penalize predicted PoP on the fixed, original scale (thresholds ***b***).

Building upon **WSR**1, we used in **WSR**2 flat-bottomed *ε*-insensitive loss functions and ***b***-transformed thresholds ***τ*** that are flexibly parameterzied and jointly learned along with other parameters from data. Furthermore, in **WSR**3 we replaced the linear decision score function *z*(·) with an RBF-kernelized one. Accordingly **WSR**2 and **WSR**3 drastically improved the performance compared to **WSR**1 for both mAUC and ordinal AUC as well as for both test sets. In particular, compared to **WSR**1, **WSR**3 improved for the ClinVar test set by 14% (14%) and did so for the CAGI set by 16% (14%) in mAUC (ordinal mAUC). Compared to the baseline **MLR** widely used in the Challenge, **WSR**3 improved for the ClinVar test set by 15% (19%) and did so for the CAGI set by 17% (17%) in mAUC (ordinal mAUC).

We found similar trends in the case of merged two-class evaluations (Class 1&2 v.s. Class 4&5). Compared to **MLR** for the CAGI set, **WSR**3 increased binary AUC from 72% to 97% and reduced RMSD from 0.28 to 0.16. Speaking of binary AUC, **WSR**3, using only 9 features, did on par with the best performer for the Challenge, LEAP (Lai et al.) even though LEAP used many more features and patient information (Cline et al., to appear).

### 3.3 Mutation-Specific Interpretation of Machine Learning Models

We went on to interpret our weakly supervised regressors, in particular **WSR**2 whose decision score function *z*(·) is linear in ***x*** and model is easily interpretable. As variants can impact disease severity through different molecular and cellular mechanisms, we ranked for each pathogenic or likely pathogenic variant their individual important features (P-value below 0.01) by P-values found by our model-interpretation procedure in Sec. 2.4.7.

As shown in Table 8, the more interpretable **WSR**2 correctly predicted 9 of 16 pathogenic or likely pathogenic to be so compared to **WSR**3 that correctly did so for 14 of those 16, which shows the trade-off between interpretability and accuracy. We focused on the 6 correctly predicted pathogenic variants and found that the most important features (and likely most probable molecular mechanisms) are related to 2 - allosteric site (BRCA1 M18T), 8 metal binding (BRCA1 C44F and C47Y), 5 - stability and conformational flexibility (BRCA1 S1715N), and 3 - catalytic site (BRCA2 R2659G and N3124). More detailed results on mutation-specific mechanistic interpretation can be found in Sec. 5 in the Supporting Information.

**TABLE 8.**
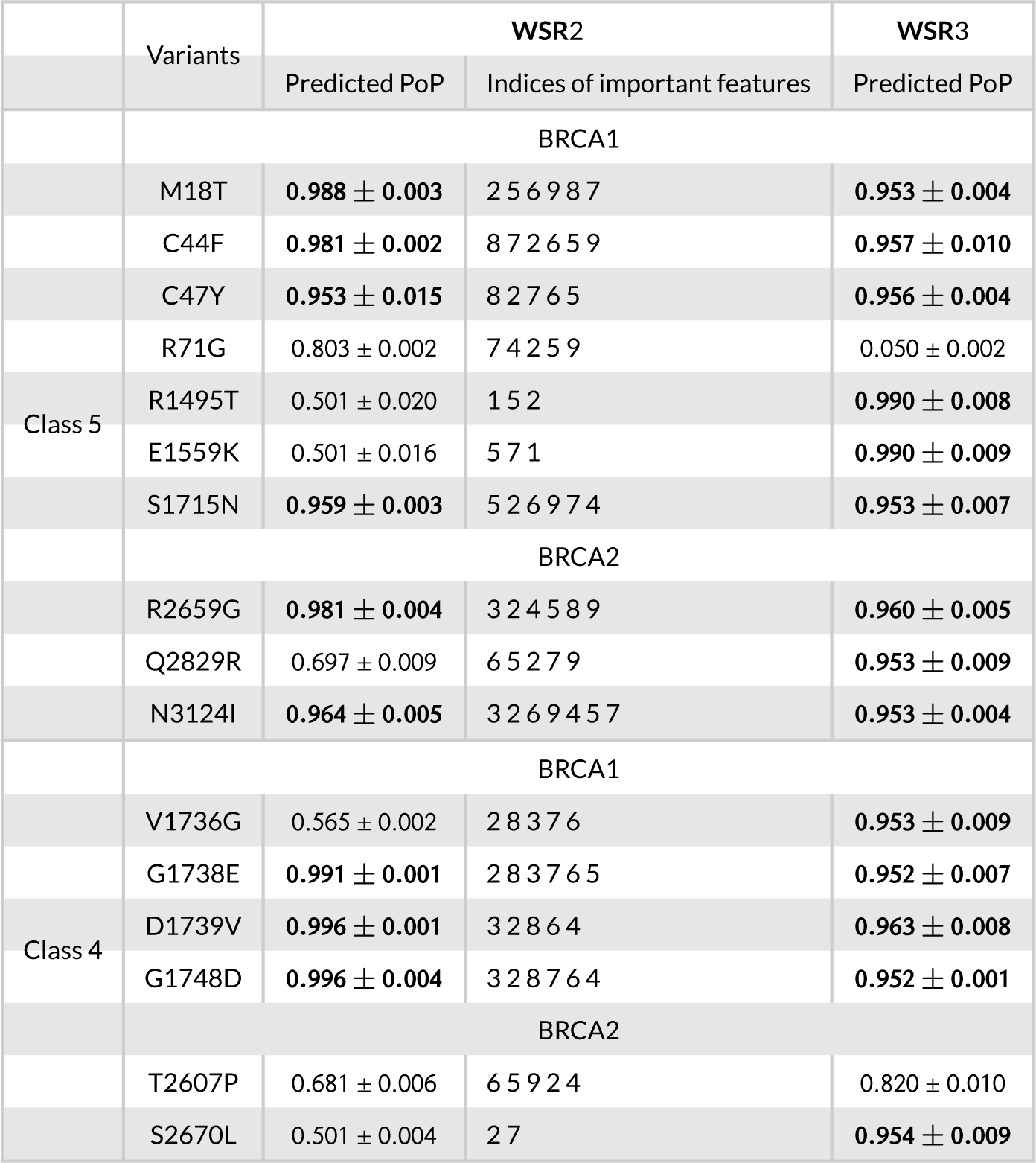
Summarized results of WSR2 model interpretation for 16 pathogenic and likely pathogenic BRCA variants. Important features are retained with a P-value cutoff of 1E-02 and ranked from left to right in increasing P-values. PoP predictions of **WSR**3, more accurate yet less interpretable, are also reported.

As will be shown next, four of these six variants with predicted mechanisms reside in available 3D protein structures and can be structurally modeled. The predicted molecular mechanisms by which mutations might confer clinical significance were thus verified to be metal binding for C44F and C47Y, stability for S1715N, and protein binding (feature 7, a top but not the first-ranked feature) for M18T.

### 3.4 Structural Interpretation of Some BRCA1 Pathogenic Variants

BRCA1 is known to be required in several cellular processes including transcription, cell- cycle check point control, DNA damage repair and control of centrosome number (Kais et al., 2012). Five pathogenic variants of BRCA1 occur at sites with available protein structure data (M18T, C44F, C47Y, and R71G in the RING domain and S1715N in the BRCT domain). In addition biological experiments have shown that M18T, C44F and C47Y are deleterious for both homologous recombination process and centrosome number, but R71G, being similar to the wild type, is deleterious for neither (Kais et al., 2012).

We performed structural modeling and energetic analysis for these variants to assess their impacts on protein folding stability and binding affinity, features found in the **WSR**2 model to be the most important features for correctly predicting pathogenicity of these variants.

#### 3.4.1 Structural modeling reproduces destabilization effects of M1775R

To validate our structural modeling protocol first, we first tried to replicate the known structure (PDB ID: 1N5O) of a pathogenic BRCA1 mutant, M1775R, by re-designing a wild-type (WT) structure (PDB ID: 1JNX). The results in **SI** Figure S1 displayed an accurate replication of mutant’s ground-truth structure. Moreover, our structural modeling provided the agreement that M1775R leads to conformational instability, a causal mechanism for its pathogenicity (Williams and Glover, 2003).

#### 3.4.2 Disrupted metal binding: C44F and C47Y

Highly conserved C44 and C47 interact with zinc ions that coordinate the stability of the RING domain (Ransburgh et al., 2010). Mutating the cystine residues would disrupt the strongly favorable sulfur-zinc interaction, which has been shown in our structural modeling as well. For both of these mutation zinc-coordinated stability has been disrupted and electrostatics is a main, disrupted molecular force based on energy decomposition (**SI** Fig. S2). Consistently shown in Table 8, feature 8 (Metal Binding) was deemed significant and ranked the first for both C44F and C47Y, which echos our structural interpretation.

#### 3.4.3 Disrupted protein binding: M18T

Heterodimerization of the RING domains of BRCA1 and BARD1 comprise an E3 ubiquitin ligase. The stability of the heterodimer is crucial for the stability of the full-length BRCA1. Mutants that do not dimerize result in defects in HDR and loss-of-tumor suppression (Starita et al., 2015). Residue M18 is at BRCA1’s interface with BARD1 and the mutation M18T is very likely to disrupt the binding with BARD1 (Morris et al., 2002). Based on structural modeling, we found that M18 of BRCA1 WT and M104 of BARD1 form a intermolecular sulfur-oxygen interaction which is known to be important for protein-protein binding (Zhang et al., 2015). We also found from structural modeling that this interaction is disrupted upon mutation M18T, which is shown in Figure 2. From binding energy decomposition (**SI** Fig. S2), electrostatics is the main reason for the binding disruption. Consistently shown in Table 8, feature 7 (Macromelecular Binding) was deemed significant (although not ranked the first) for M18T, which agrees with our structural interpretation.

**FIGURE 2.**
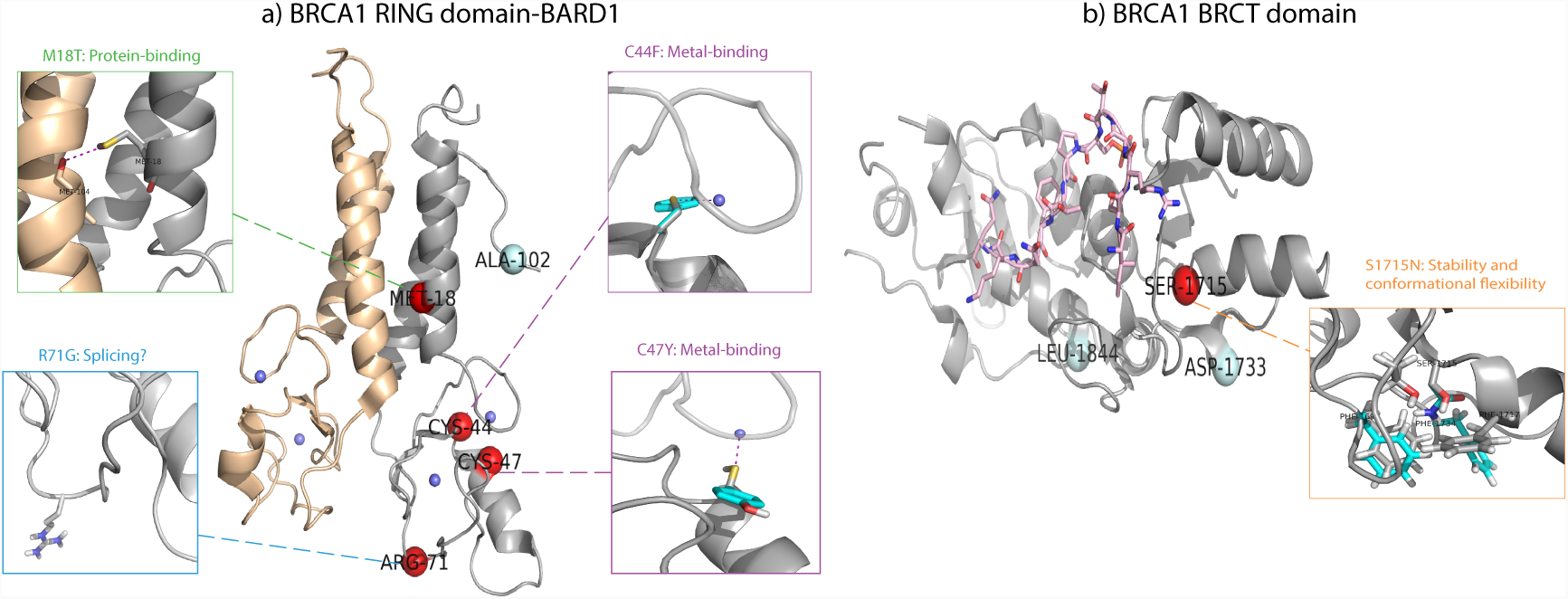
Structural interpretation of pathogenicity mechanisms for several BRCA1 variations at structurally-available RING and BRCT domains. Pathogenic (Class 5) and benign (Class 1) variation sites are shown in red and pale cyan spheres. Zoomed-in illustrations of molecular mechanisms have been shown for individual variants in smaller side boxes, where crystal wild-type residues are in gray sticks and modeled mutant residues are in cyan sticks. a) RING domain complex of BRCA1-BARD1 in PDB structure 1JM7 where RING domain of BRCA1 is shown in gray cartoon, BARD1 wheat caroon, and Zn^2+^ ions small blue sphere. b) BRCT domain of BRCA1 interacting with Bach1 helicase in PDB structure 1T29 PDB where BRCT is shown in grey cartoon and Bach1 helicase in pink sticks.

#### 3.4.4 Aberrant splicing: R71G

The mutation R71G was found to affect the splicing process rather than homologous recombination process and the centrosome number (Vega et al., 2001; Kais et al., 2012). The aberrant splicing of BRCA1 mRNA would result in premature translation and truncated proteins, which is beyond the capability of structural modeling. Indeed, the mutation was not found to destabilize BRCA1 (**SI** Fig. S2). Interestingly, the top-performing LEAP team found that splicing information, such as the distance to the nearest splice site, was also important for correctly annotating the clinical significance of some variants (Cline et al., to appear).

#### 3.4.5 Decreased stability: S1715N

The BRCT domain of BRCA1 displays an intrinsic transactivation activity. S1715 is an evolutionarily conserved residue and pathogenic mutations in the BRCA domain including S1715N have shown the loss of such activity in yeast and mammalian cells (Vallon-Christersson et al., 2001). S1715N has been shown unable to complement BRCA1 deficiency for homologous recombination and leading to high instability (Petitalot et al., 2019). Our structural modeling confirms that S1715N is structurally destabilizing (**SI** Fig. S2). Specifically, van der Waals clashes was found the main reason for the destabilization effect, as shown in **SI** Fig. S2. Of course, the extent of such clashes could be allieviated by improving the modeling of structure flexibility. Consistently show in Table 8, feature 5 (Stability and Conformational Flexibility) was ranked the most important for S1715N, echoing our structural interpretation.

To summarize, we have found for these pathogenic BRCA1 variants the consensus of our machine learning-based and structure modeling-based mechanistic interpretations. Although we do not have structure data to start with for some mutation sites, our machine learning-based interpretations could generate actionable hypotheses of the causal mechanisms that can be tested experimentally, potentially suggesting therapeutic candidates accordingly.

## 4 CONCLUSION

Starting with interpretable and actionable features that capture molecular impacts of genetic variants and expert-curated albeit inexact labels that only annotate variants using their ranges in the probability of pathogenicity (PoP), we have developed novel weakly-supervised regression models that can directly predict the probability of pathogenicity. By considering the order among pathogenicity classes, penalizing PoP prediction with novel loss functions, and embedding the original feature space into a kernel space, our weakly supervised regressor 3 – **WSR**3 – has significantly improved the predictive performance for a CAGI challenge set compared to a representative multi-class classification model: binary AUC increased from 0.72 to 0.97 and ordinal multi-class AUC increased from 0.64 to 0.80. **WSR**3 even improved the predictive performances compared to a representative ordinal regression model **CLM**(again, a multi-class classifier) that respects class order. Note that our pathogenicity predictors are not gene-specific but gene type-specific, which could access more variants data and more advanced machine learning models.

We further developed methods to interpret our weakly-supervised regressor by assessing the statistical significance of feature importance in supporting individual model predictions. We identified and ranked important features (each corresponding to a mechanism of molecular impacts upon amino-acid substitution) for newly annotated or updated, pathogenic or likely pathogenic BRCA variants. Our structural modeling of mutational effects on protein folding stability and binding affinity has corroborated the machine learning-predicted molecular mechanisms by which genetic variants lead to diseases. Namely, validated in structural modeling, metal binding for BRCA1 C44F and C47Y was predicted by our machine learning model interpretation as the most important feature for the pathogenic calling of the variants. So was stability for S1715N. And protein binding was predicted to be a statistically significant but not the first-ranked feature for M18T. These promising results indicate that these models could generate experimentally-testable mechanistic hypotheses and lead to therapeutic candidates accordingly.

We only used nine features out of over 50 property probabilities from MutPred2 and could have achieved even better performances with more features although highly dependent or not directly causal features could hurt model interpretability. One limitation about the current feature set though is the entire focus on molecular impacts of genetic variation without the consideration of cellular contexts or systems-level impacts. On one hand, some variants significantly impacting protein functions may not lead to clinical significance. On the other hand, splicing information for some missense variants is found important for their pathogenicity prediction(Cline et al., to appear). Therefore, it would be of great value to include endophenotypes (Masica and Karchin, 2016) encoding causal mechanisms across hierarchical subsystems (Yu et al., 2016) at various biological scales. Although endophenotype predictors are not adequate for the purpose yet, increasingly available big data and empowering artificial intelligence methods (Ma et al., 2018) are stimulating their development and making the goal of precision medicine more attainable than ever.

## ACKNOWLEDGMENTS

We thank the CAGI organizers, data providers, challenge assessors, and fellow participants for the insightful challenges, data, analysis, and tools (in particular, MutPred2) which made this study possible. We also thank Dr. Vikas Rao Pejaver for the help on generating Mutpred2 features. Part of the computing support was provided by the Texas A&M University High Performance Research Computing.

## Abbreviations

CAGI: Critical Assessment of Genome Interpretation
ENIGMA: Evidence-based Network for the Interpretation of Germline Mutant Alleles
PoP: Probability of Pathogenicity
WSR: Weekly Supervised Regression
MLR: Multi-class Logistic Regression
CLM: Cumulative Logit Model
AUC: Area Under the Curve

