## Supporting Information for "Predicting Pathogenicity of Missense Variants withWeakly Supervised Regression"

### Predicting Pathogenicity of Missense Variants with Weakly Supervised Regression (SI)

#### 1 Data Collection

The complete pipeline of collecting data for BRCA1/2 or other genes is explained below.

##### 1.1 Generation of gene candidates

We used the STRING database [Szklarczyk et al., 2017] to find other genes whose encoded proteins have close interaction with BRCA1 and BRCA2 proteins. We chose multiple proteins mode only changing the follow setting parameters: 1) set first shell under 'max number of interactors to show' to 150; 2) set the custom value of 'minimum required interaction score' to 0.95 which we assumed as a reasonable lower bound. Such setting gave us 74 genes including BRCA1 and BRCA2.

##### 1.2 Tumor suppressor genes filtering

Then we selected tumor suppressor genes within these 74 genes according to the annotation from OncoKB [Chakravarty et al., 2017]. We ended up with 23 tumor suppressor genes including BRCA1/2.

##### 1.3 Retrieving variants and label data

We queried ClinVar dataset with filters and values shown below. One query command example for BRCA1 variants with review status greater or equal to 2 stars is: *((BRCA1[Gene Name]) AND ("1990/01/01"[Last interpreted] : "2017/06/29"[Last interpreted]) AND ( ( ("clinsig benign"[Properties]) OR ("clinsig likely benign"[Properties]) OR ("clinsig vus"[Properties]) OR ("clinsig likely pathogenic"[Properties]) OR ("clinsig pathogenic"[Properties]) ) AND ( ("practice guideline"[Review status]) OR ("reviewed by expert panel"[Review status]) OR ("criteria provided, multiple submitters, no conflicts"[Review status]) ) AND ("missense variant"[molecular consequence])))*. To search variants with review status greater or equal to 3 stars, remove *OR ("criteria provided, multiple submitters, no conflicts"[Review status])* part in the query command. The query filters used are summarized as follows:

| Field | Value |
|---|---|
| Gene name | e.g. BRCA1 |
| Last interpreted | 1990/01/01 - 2017/06/29 |
| Properties | clinsig benign; clinsig likely benign; clinsig vus;<br>clinsig likely pathogenic; clinsig pathogenic |
| Review status | practice guideline(4 stars); reviewed by expert panel(3 stars);<br>criteria provided, multiple submitters, no conflicts(2 stars) |
| Molecular consequence | missense variant |

Table S1: ClinVar query filters and values

We further extracted the 'clinical significance' column as labels from the variant records downloaded from ClinVar. There are two cases to be taken care of. First, some variants are interpreted with multiple pathogenicity classes, such as *benign/likely benign* or *pathogenic/likely pathogenic*. In this case we adopted the more conservative labels, such as likely benign for benign/likely benign and pathogenic for pathogenic/likely pathogenic. Second, some variant records may correspond to different single nucleotide variants (SNVs) but the same single amino-acid variant (SAV). This case involved only scores of variants that were not included in our training data.

4 of 23 genes have no variant annotation in ClinVar and the final data set contains those for 19 genes (including BRCA1/2).

##### 1.4 Generating input features

We collected protein sequence data from Uniprot[Bateman et al., 2017] and generated input FASTA files for Mutpred2 [Pejaver et al., 2017] following its requirement. Molecular properties are organized hierarchically forming a tree structure. We extracted these 9 third level properties for each variant and assembled them into a feature file for each protein. We set 0 for properties with missing values for one variant.

#### 2 Data Statistics

The statistics for the raw data sets **G2** and **G3** (at least 2 stars or 3 stars in ClinVar 'review status') before being split into training and held-out testing sets is listed in Table S3 and Table S2, respectively. The data statistics of the CAGI test set is shown in Table S4.

| Genes | 1-Benign | 2-Likely Benign | 3-Uncertain Significance | 4-Likely Pathogenic | 5-Pathogenic | Total |
|---|---|---|---|---|---|---|
| BRCA1 | 87 | 0 | 0 | 0 | 21 | 108 |
| BRCA2 | 84 | 0 | 0 | 0 | 9 | 93 |
| Sub Total (BRCA1/2 only) | 171 (85.07%) | 0 (0.0%) | 0 (0.0%) | 0 (0.0%) | 30 (14.93%) | 201 |
| MLH1 | 17 | 9 | 161 | 18 | 55 | 260 |
| Other genes | 0 | 0 | 0 | 0 | 0 | 0 |

Table S2: Statistics for all genes in the **G3** data set

#### 3 Theoretical Results for Kernelizing Weakly Supervised Regression

**Mathematical Notation**: For two vectors $\boldsymbol{a}$ and $\boldsymbol{b}$, $\{\boldsymbol{a}, \boldsymbol{b}\}$ means the concatenated vector of $\boldsymbol{a}$ and $\boldsymbol{b}$.

**Lemma 1 (Representer Theorem).** *Let $\mathcal{X}$ be a nonempty set and $\kappa$ a positive-definite real-valued kernel on $\mathcal{X}\times\mathcal{X}$ with corresponding reproducing kernel Hilbert space $H_\kappa$. Given n training samples $(\boldsymbol{x}_1, y_1), \ldots, (\boldsymbol{x}_n, y_n) \in \mathcal{X} \times R$, a non-decreasing real-valued function $g\colon [0,\infty) \to R$, and a function $L\colon (\mathcal{X} \times R^2)^n \to R$, then for any $f^* \in H_\kappa$ satisfying*

$$f^* = \underset{f\in H_\kappa}{\operatorname{argmin}} \left\{L\left((f(\boldsymbol{x}_1), y_1), ..., (f(\boldsymbol{x}_n), y_n)\right) + g\left(||f||_{H_\kappa}\right)\right\}, \tag{1}$$

*$f^*$ admits a representation of the form:*

$$f^*(\cdot) = \sum_{i=1}^{n} \alpha_i \kappa(\cdot, \boldsymbol{x}_i) \tag{2}$$

**Theorem 1.** *Let $(\boldsymbol{x}_1, y_1), (\boldsymbol{x}_2, y_2), ..., (\boldsymbol{x}_n, y_n)$ be the training samples. The optimal parameters of the following optimization problem*

$$\begin{aligned} \min_{\boldsymbol{w},\boldsymbol{\tau}} \quad & f(\boldsymbol{w}, \boldsymbol{\tau}) = \frac{1}{n}\sum_{i}^{n} \ell(\boldsymbol{w}^T\boldsymbol{x}_i, \boldsymbol{\tau}, y_i) + \lambda||\boldsymbol{w}||_2^2 \\ s.\ t. \quad & g(\boldsymbol{\tau}) \geq 0 \end{aligned} \tag{3}$$

| Genes | 1-Benign | 2-Likely Benign | 3-Uncertain Significance | 4-Likely Pathogenic | 5-Pathogenic | Total |
|---|---|---|---|---|---|---|
| BRCA1 | 87 | 2 | 153 | 2 | 28 | 272 |
| BRCA2 | 84 | 3 | 330 | 1 | 9 | 427 |
| MLH1 | 17 | 9 | 205 | 20 | 55 | 306 |
| BRIP1 | 0 | 0 | 55 | 0 | 0 | 55 |
| BARD1 | 0 | 0 | 48 | 0 | 0 | 48 |
| PALB2 | 0 | 1 | 67 | 0 | 0 | 68 |
| RAD50 | 0 | 0 | 47 | 0 | 0 | 47 |
| NBN | 0 | 0 | 35 | 0 | 0 | 35 |
| TP53 | 0 | 0 | 29 | 3 | 4 | 36 |
| ATM | 0 | 0 | 174 | 2 | 1 | 177 |
| FANCD2 | 0 | 0 | 1 | 0 | 0 | 1 |
| MRE11A (MRE11) | 0 | 0 | 19 | 0 | 0 | 19 |
| RAD51C | 0 | 0 | 13 | 0 | 0 | 13 |
| ATR | 0 | 7 | 2 | 0 | 0 | 9 |
| BLM | 0 | 2 | 5 | 0 | 0 | 7 |
| XRCC2 | 0 | 0 | 6 | 0 | 0 | 6 |
| FAM175A (ABRAXAS1) | 0 | 3 | 3 | 0 | 0 | 6 |
| CHEK2 | 0 | 0 | 47 | 0 | 0 | 47 |
| FANCA | 3 | 6 | 5 | 1 | 0 | 15 |
| Total | 191 (11.98%) | 33 (2.07%) | 1244 (78.04%) | 29 (1.82%) | 97 (6.09%) | 1594 |

Table S3: Statistics for all genes in the **G2** data set

| Genes | 1-Benign | 2-Likely Benign | 3-Uncertain Significance | 4-Likely Pathogenic | 5-Pathogenic | Total |
|---|---|---|---|---|---|---|
| BRCA1 | 31 | 100 | 2 | 4 | 7 | 144 |
| BRCA2 | 31 | 136 | 2 | 2 | 3 | 174 |
| Total | 62 (19.50%) | 236 (74.21%) | 4 (1.26%) | 6 (1.89%) | 10 (3.14%) | 318 |

Table S4: Statistics for the CAGI test set used in the ENIGMA Challenge.

*could be written as:*

$$\{\boldsymbol{w}^*, \boldsymbol{\tau}^*\} = \{\sum_{i=1}^{n} \beta_i \boldsymbol{x}_i^T, \boldsymbol{\tau}^*\} \tag{4}$$

*where* $\boldsymbol{\beta} = (\beta_1, \beta_2, ..., \beta_n)$ *are the new parameters. And the optimization problem above could be kernelized as:*

$$\begin{aligned} \min_{\boldsymbol{\beta},\boldsymbol{\tau}} \quad & \tilde{f}(\boldsymbol{\beta}, \boldsymbol{\tau}) = \frac{1}{n}\sum_{i}^{n} \ell(\sum_{j} \beta_j \kappa(\boldsymbol{x}_i, \boldsymbol{x}_j), \boldsymbol{\tau}, y_i) + \lambda \boldsymbol{\beta}^T \boldsymbol{K} \boldsymbol{\beta} \\ s.\ t. \quad & g(\boldsymbol{\tau}) \geq 0 \end{aligned} \tag{5}$$

*where* $\kappa(\cdot,\cdot)$ *is a positive-defined real-valued kernel, and* $\boldsymbol{K}$ *is the kernel matrix where* $[\boldsymbol{K}]_{ij} = \kappa(\boldsymbol{x}_i, \boldsymbol{x}_j)$.

*Proof.* Without the loss of generalizability, we let $\{\boldsymbol{w}^*, \boldsymbol{\tau}^*\}$ be one of the optimal parameters of the above optimization problem. We then fix $\boldsymbol{\tau} = \boldsymbol{\tau}^*$ as constants. Since the constraints in the above optimization problem do not contain $\boldsymbol{w}$ and are satisfied when $\boldsymbol{\tau} = \boldsymbol{\tau}^*$, we can drop the constraints and solve the following unconstrained optimization problem:

$$\min_{\boldsymbol{w}} \quad f(\boldsymbol{w}, \boldsymbol{\tau} = \boldsymbol{\tau}^*) \tag{6}$$

Based on **Lemma 1**, every optimal solution: $\boldsymbol{w}^{*\prime} = \underset{\boldsymbol{w}}{\operatorname{argmin}} f(\boldsymbol{w}, \boldsymbol{\tau} = \boldsymbol{\tau}^*)$ can be written as the linear combination of all training samples: $\boldsymbol{w}^{*\prime} = \sum_{i=1}^{n} \beta_i \boldsymbol{x}_i^T$.

Since $(\boldsymbol{w}^*, \boldsymbol{\tau}^*) = \underset{\boldsymbol{w},\boldsymbol{\tau}}{\operatorname{argmin}}\, f(\boldsymbol{w}, \boldsymbol{\tau})$, $\boldsymbol{w}^*$ must also be the optimal parameters of $\min_{\boldsymbol{w}} f(\boldsymbol{w}, \boldsymbol{\tau} = \boldsymbol{\tau}^*)$. Therefore, $\boldsymbol{w}^*$ could also be written as the linear combination of all training samples as Eq 4. We then put Eq 4 into the objective function of the original optimization problem and get:

$$\tilde{f}(\boldsymbol{w}, \boldsymbol{\tau}) = \frac{1}{n} \sum_{i}^{n} \ell(\sum_{j} \beta_j \boldsymbol{x}_j^T \boldsymbol{x}_i, \boldsymbol{\tau}, y_i) + \lambda \sum_{i} \sum_{j} \beta_i \beta_j \boldsymbol{x}_i^T \boldsymbol{x}_j \tag{7}$$

We replace the inner product $\boldsymbol{x}_i^T \boldsymbol{x}_j$ with the kernel function $k(\boldsymbol{x}_i, \boldsymbol{x}_j)$ in Eq 7, and put the constraint back. We finally get the exact format of the optimization problem shown in Eq 5, which concludes the proof.

#### 4 Optimization Results for WSR2 and WSR3

Recall in the main text, we used 100 randomly distributed starting points to solve nonconvex optimziation problems through BFGS (unconstrained) or L-BFGS-B (box-constrained). We analyze the optimization results for both the linear and RBF kernelized model – **WSR**. Let $f_i$ be the final objective value after the $i$th optimization trajectory, $f_{min} = \min_i \{f_i\}$ and $\sigma_i = \frac{f_i - f_{min}}{f_{min}}$ with $1 \leq i \leq 100$. For simplicity, we only assess the 100 trajectories' optimization results for the optimal hyper-parameters reported in Table(S5).

We withheld one of the five folds of training data one by one. Each time we end up with 4-fold cross validation for the optimal hyperparameters. Once the optimal hyperparameters are found, we fix the hyper-parameters values and train the parameters using all 4 folds. Therefore, for each of the five such processes, we uniformly sample 100 initial starting points for $(\boldsymbol{w}, \boldsymbol{\tau})$ for parameter training. After all 500 trajectories of optimization, we calculate the frequency of $\sigma_i$ that is below a given threshold. We can see from Table S6, Nearly 70% of $\sigma_i$s are below 0.1% in both linear and RBF kernelized models, which means around 70 out of 100 initial points will lead to the same optimized performance.

We further study the locations of optima found. We let $\boldsymbol{x}_{min}$ be the minimum corresponding to $f_{min}$ and $\epsilon_i = \frac{||\boldsymbol{x}_i - \boldsymbol{x}_{min}||}{||\boldsymbol{x}_{min}||}$ with $1 \leq i \leq 100$. Considering the precision level used in L-BFGS-B, we think if $\epsilon_i < 0.1\%$ then $\boldsymbol{x}_i$ and $\boldsymbol{x}_{min}$ are numerically equal. After optimization, **63%** and **60%** of $\epsilon_i$s are found to be lower than 0.1% in linear and RBF kernelized models, respectively. This indicates that more than 60 out of 100 initial points will be optimized not only to reach the same optimal value but also the same optimum. This result shows that a wide attraction basin around the found optimum may exist in the landscape of the loss functions involved.

| Hyperparameters | $\lambda$ | $\alpha_1$ | $\alpha_2$ | $\alpha_3$ | $\alpha_4$ | $\alpha_5$ | $\gamma$ |
|---|---|---|---|---|---|---|---|
| **WSR**2 | 1.73 | 1.41 | 1.0 | 0.71 | 1.0 | 1.41 | N.A. |
| **WSR**3 | 0.76 | 1.41 | 1.0 | 0.71 | 1.0 | 1.41 | 0.50 |

Table S5: Optimal hyperparameters after training in **WSR**2&3 through **G2** with all genes.

| Threshold | 5% | 1% | 0.1% | 0.01% |
|---|---|---|---|---|
| Frq.(Per.) in **WSR**2 | 471(94%) | 451(90%) | 375(75%) | 344(69%) |
| Frq.(Per.) in **WSR**3 | 465(93%) | 433(87%) | 347(69%) | 315(63%) |

Table S6: The percentage(frequency) of $\sigma_i$s over 500 under a given threshold in **WSR**2&3.

#### 5 Mutation-specific Machine Learning Model Interpretation

The mechanistic interpretations of pathogenic or likely pathogenic BRCA variants in the ENIGMA Challenge (CAGI test set) are shown from **Table** S7. "PoP" stands for the probability of pathogenicity.

| Variant: **M18T**, PoP=0.988 $\pm$ 0.003 | | | |
|---|---|---|---|
| Feature Index | Feature name | P-value | T-statistic |
| 2 | Allosteric site | 2.94E-167 | 82.53 |
| 5 | Stability and conformational flexibility | 7.48E-92 | 34.96 |
| 6 | Special structural signatures | 1.08E-79 | 29.95 |
| 9 | PTM site | 2.09E-43 | 17.38 |
| 8 | Metal binding | 2.75E-07 | 5.16 |
| 7 | Macromolecular binding | 2.25E-03 | 2.02 |
| 4 | Secondary structure | 9.09E-01 | -1.34 |
| 3 | Catalytic site | 1.00E+00 | -3.60 |
| 1 | Relative solvent accessibility | 1.00E+00 | -27.88 |

| Variant: **C44F**, PoP=0.981 $\pm$ 0.002 | | | |
|---|---|---|---|
| Feature Index | Feature name | P-value | T-statistic |
| 8 | Metal binding | 2.68E-160 | 76.52 |
| 7 | Macromolecular binding | 1.37E-118 | 48.05 |
| 2 | Allosteric site | 1.72E-99 | 38.38 |
| 6 | Special structural signatures | 1.08E-79 | 29.95 |
| 5 | Stability and conformational flexibility | 3.04E-62 | 23.55 |
| 9 | PTM site | 4.97E-20 | 10.03 |
| 1 | Relative solvent accessibility | 3.94E-05 | 4.02 |
| 4 | Secondary structure | 9.09E-01 | -1.34 |
| 3 | Catalytic site | 1.00E+00 | -3.60 |

| Variant: **C47Y**, PoP=0.953 $\pm$ 0.015 | | | |
|---|---|---|---|
| Feature Index | Feature name | P-value | T-statistic |
| 8 | Metal binding | 7.49E-147 | 66.02 |
| 2 | Allosteric site | 1.72E-99 | 38.38 |
| 7 | Macromolecular binding | 2.56E-92 | 35.16 |
| 6 | Special structural signatures | 5.82E-86 | 32.47 |
| 5 | Stability and conformational flexibility | 3.63E-10 | 6.45 |
| 9 | PTM site | 2.00E-01 | 0.84 |
| 1 | Relative solvent accessibility | 4.86E-01 | 0.04 |
| 3 | Catalytic site | 1.00E+00 | -3.60 |
| 4 | Secondary structure | 1.00E+00 | -6.10 |

| Variant: **R71G**, PoP=0.803 $\pm$ 0.002 | | | |
|---|---|---|---|
| Feature Index | Feature name | P-value | T-statistic |
| 7 | Macromolecular binding | 1.76E-58 | 22.27 |
| 4 | Secondary structure | 4.43E-29 | 12.93 |
| 2 | Allosteric site | 9.32E-26 | 11.88 |
| 5 | Stability and conformational flexibility | 3.63E-10 | 6.45 |
| 9 | PTM site | 5.09E-06 | 4.52 |
| 6 | Special structural signatures | 1.22E-02 | 2.27 |
| 1 | Relative solvent accessibility | 4.86E-01 | 0.04 |
| 3 | Catalytic site | 1.00E+00 | -3.60 |
| 8 | Metal binding | 1.00E+00 | -7.43 |

| Variant: **R1495T**, PoP=0.501 $\pm$ 0.020 | | | |
|---|---|---|---|
| Feature Index | Feature name | P-value | T-statistic |
| 1 | Relative solvent accessibility | 3.98E-26 | 12.00 |
| 5 | Stability and conformational flexibility | 1.11E-20 | 10.25 |
| 2 | Allosteric site | 1.28E-03 | 3.05 |
| 7 | Macromolecular binding | 4.30E-01 | 0.18 |
| 4 | Secondary structure | 9.09E-01 | -1.34 |
| 3 | Catalytic site | 1.00E+00 | -3.60 |
| 6 | Special structural signatures | 1.00E+00 | -5.29 |
| 8 | Metal binding | 1.00E+00 | -7.43 |
| 9 | PTM site | 1.00E+00 | -23.04 |

Table S7: The results of one-tailed one-sample $t$-test for each pathogenic mutation based on weakly supervision regression (WSR2). T-statistic is calculated through $\frac{\bar{x}}{s/\sqrt{n}}$, where $\bar{x}$, $s$ and $n$ are the sample mean, sample standard deviation and sample size, respectively. Features are ranked by P-value.

| Variant: **E1559K**, PoP=0.501 ± 0.016 | | | |
|---|---|---|---|
| Feature Index | Feature name | P-value | T-statistic |
| 5 | Stability and conformational flexibility | 1.24E-56 | 21.65 |
| 7 | Macromolecular binding | 1.87E-35 | 14.90 |
| 1 | Relative solvent accessibility | 3.34E-14 | 8.01 |
| 3 | Catalytic site | 1.00E+00 | -3.60 |
| 4 | Secondary structure | 1.00E+00 | -6.10 |
| 8 | Metal binding | 1.00E+00 | -7.43 |
| 6 | Special structural signatures | 1.00E+00 | -7.80 |
| 2 | Allosteric site | 1.00E+00 | -14.61 |
| 9 | PTM site | 1.00E+00 | -19.37 |

| Variant: **S1715N**, PoP=0.959 ± 0.003 | | | |
|---|---|---|---|
| Feature Index | Feature name | P-value | T-statistic |
| 5 | Stability and conformational flexibility | 6.74E-78 | 29.25 |
| 2 | Allosteric site | 8.43E-54 | 20.71 |
| 6 | Special structural signatures | 2.97E-51 | 19.88 |
| 9 | PTM site | 3.40E-49 | 19.22 |
| 7 | Macromolecular binding | 1.87E-35 | 14.90 |
| 4 | Secondary structure | 3.80E-04 | 3.42 |
| 1 | Relative solvent accessibility | 4.86E-01 | 0.04 |
| 3 | Catalytic site | 1.00E+00 | -3.60 |
| 8 | Metal binding | 1.00E+00 | -5.33 |

| Variant: **R2659G**, PoP= 0.981 ± 0.004 | | | |
|---|---|---|---|
| Feature Index | Feature name | P-value | T-statistic |
| 3 | Catalytic site | 8.25E-185 | 99.75 |
| 2 | Allosteric site | 1.72E-99 | 38.38 |
| 4 | Secondary structure | 1.08E-84 | 31.95 |
| 5 | Stability and conformational flexibility | 3.63E-10 | 6.45 |
| 8 | Metal binding | 1.24E-03 | 3.06 |
| 9 | PTM site | 3.95E-03 | 2.68 |
| 7 | Macromolecular binding | 2.25E-02 | 2.02 |
| 1 | Relative solvent accessibility | 4.86E-01 | 0.04 |
| 6 | Special structural signatures | 9.97E-01 | -2.77 |

| Variant: **Q2829R**, PoP= 0.697 ± 0.009 | | | |
|---|---|---|---|
| Feature Index | Feature name | P-value | T-statistic |
| 6 | Special structural signatures | 3.74E-66 | 24.92 |
| 5 | Stability and conformational flexibility | 7.65E-51 | 19.75 |
| 2 | Allosteric site | 9.32E-26 | 11.88 |
| 7 | Macromolecular binding | 7.52E-05 | 3.86 |
| 9 | PTM site | 3.95E-03 | 2.68 |
| 1 | Relative solvent accessibility | 4.86E-01 | 0.04 |
| 8 | Metal binding | 9.99E-01 | -3.23 |
| 3 | Catalytic site | 1.00E+00 | -3.60 |
| 4 | Secondary structure | 1.00E+00 | -6.10 |

| Variant: **N3124I**, PoP= 0.964 ± 0.005 | | | |
|---|---|---|---|
| Feature Index | Feature name | P-value | T-statistic |
| 3 | Catalytic site | 1.44E-212 | 134.20 |
| 2 | Allosteric site | 1.72E-99 | 38.38 |
| 6 | Special structural signatures | 4.29E-73 | 27.43 |
| 9 | PTM site | 1.38E-31 | 13.71 |
| 4 | Secondary structure | 1.22E-14 | 8.17 |
| 5 | Stability and conformational flexibility | 3.63E-10 | 6.45 |
| 7 | Macromolecular binding | 7.52E-05 | 3.86 |
| 8 | Metal binding | 9.99E-01 | -3.23 |
| 1 | Relative solvent accessibility | 1.00E+00 | -23.89 |

Table S7: (Continued) The results of one-tailed one-sample $t$-test for each pathogenic mutation based on weakly supervision regression (WSR2). T-statistic is calculated through $\frac{\bar{x}}{s/\sqrt{n}}$, where $\bar{x}$, $s$ and $n$ are the sample mean, sample standard deviation and sample size, respectively. Features are ranked by P-value.

### 6 Replicating the crystal structure of M1775R mutants by redesigning the WT

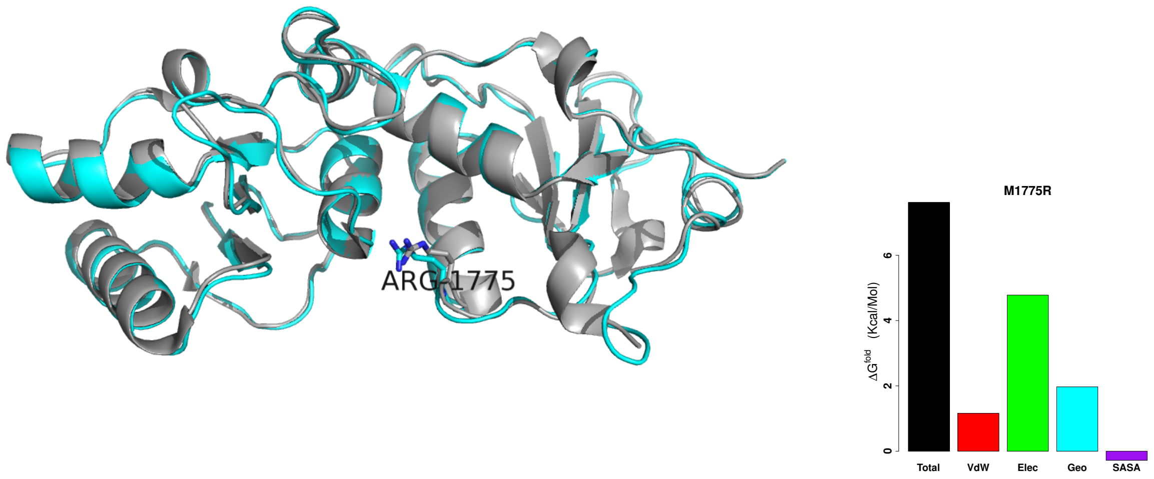


Fig. S1: Replicating the crystal structure of M1775R mutants (PDB accession code:1N5O) from the WT (PDB accession code:1JNX) and its energy decomposition which shows that electrostatic is the main reason for disrupting the stability of BRCT domain.

### 7 Structural Modeling of Newly Annotated Pathogenic BRCA1 Variants

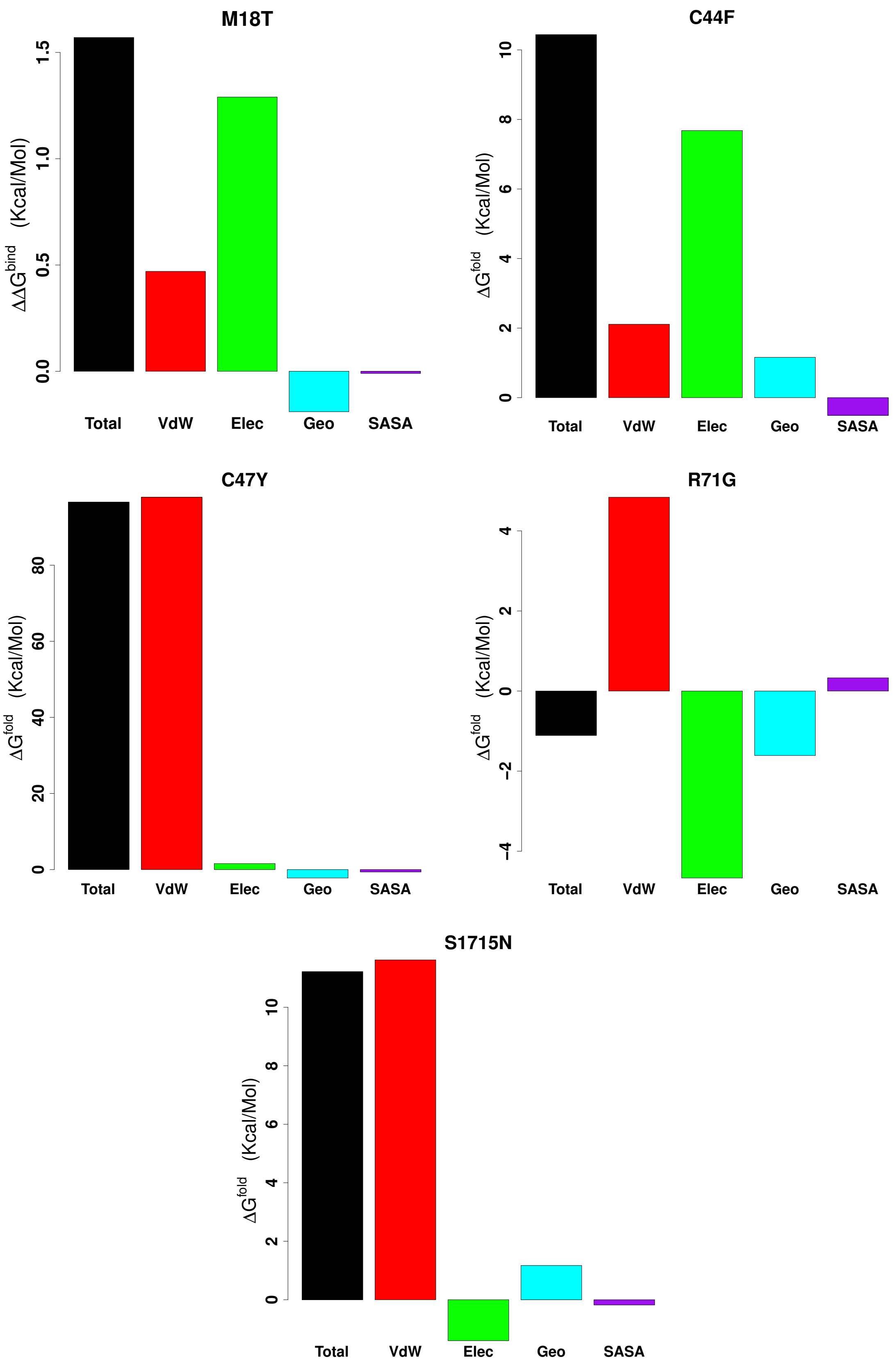


Fig. S2: Energy decomposition for mutants based on structure-based redesign by iCFN [Karimi and Shen, 2018].

### Bibliography

Alex Bateman, Maria Jesus Martin, Claire ODonovan, Michele Magrane, Emanuele Alpi, Ricardo Antunes, Benoit Bely, Mark Bingley, Carlos Bonilla, Ramona Britto, Borisas Bursteinas, Hema Bye-A-Jee, Andrew Cowley, Alan Da Silva, Maurizio De Giorgi, Tunca Dogan, Francesco Fazzini, Leyla Garcia Castro, Luis Figueira, Penelope Garmiri, George Georghiou, Daniel Gonzalez, Emma Hatton-Ellis, Weizhong Li, Wudong Liu, Rodrigo Lopez, Jie Luo, Yvonne Lussi, Alistair MacDougall, Andrew Nightingale, Barbara Palka, Klemens Pichler, Diego Poggioli, Sangya Pundir, Luis Pureza, Guoying Qi, Alexandre Renaux, Steven Rosanoff, Rabie Saidi, Tony Sawford, Aleksandra Shypitsyna, Elena Speretta, Edward Turner, Nidhi Tyagi, Vladimir Volynkin, Tony Wardell, Kate Warner, Xavier Watkins, Rossana Zaru, Hermann Zellner, Ioannis Xenarios, Lydie Bougueleret, Alan Bridge, Sylvain Poux, Nicole Redaschi, Lucila Aimo, Ghislaine Argoud-Puy, Andrea Auchincloss, Kristian Axelsen, Parit Bansal, Delphine Baratin, Marie-Claude Blatter, Brigitte Boeckmann, Jerven Bolleman, Emmanuel Boutet, Lionel Breuza, Cristina Casal-Casas, Edouard de Castro, Elisabeth Coudert, Beatrice Cuche, Mikael Doche, Dolnide Dornevil, Severine Duvaud, Anne Estreicher, Livia Famiglietti, Marc Feuermann, Elisabeth Gasteiger, Sebastien Gehant, Vivienne Gerritsen, Arnaud Gos, Nadine Gruaz-Gumowski, Ursula Hinz, Chantal Hulo, Florence Jungo, Guillaume Keller, Vicente Lara, Philippe Lemercier, Damien Lieberherr, Thierry Lombardot, Xavier Martin, Patrick Masson, Anne Morgat, Teresa Neto, Nevila Nouspikel, Salvo Paesano, Ivo Pedruzzi, Sandrine Pilbout, Monica Pozzato, Manuela Pruess, Catherine Rivoire, Bernd Roechert, Michel Schneider, Christian Sigrist, Karin Sonesson, Sylvie Staehli, Andre Stutz, Shyamala Sundaram, Michael Tognolli, Laure Verbregue, Anne-Lise Veuthey, Cathy H. Wu, Cecilia N. Arighi, Leslie Arminski, Chuming Chen, Yongxing Chen, John S. Garavelli, Hongzhan Huang, Kati Laiho, Peter McGarvey, Darren A. Natale, Karen Ross, C. R. Vinayaka, Qinghua Wang, Yuqi Wang, Lai-Su Yeh, and Jian Zhang. UniProt: the universal protein knowledgebase. *Nucleic Acids Research*, 45(D1):D158–D169, January 2017. ISSN 0305-1048. https://doi.org/10.1093/nar/gkw1099. URL https://academic.oup.com/nar/article/45/D1/D158/2605721.

Debyani Chakravarty, Jianjiong Gao, Sarah Phillips, Ritika Kundra, Hongxin Zhang, Jiaojiao Wang, Julia E. Rudolph, Rona Yaeger, Tara Soumerai, Moriah H. Nissan, Matthew T. Chang, Sarat Chandarlapaty, Tiffany A. Traina, Paul K. Paik, Alan L. Ho, Feras M. Hantash, Andrew Grupe, Shrujal S. Baxi, Margaret K. Callahan, Alexandra Snyder, Ping Chi, Daniel C. Danila, Mrinal Gounder, James J. Harding, Matthew D. Hellmann, Gopa Iyer, Yelena Y. Janjigian, Thomas Kaley, Douglas A. Levine, Maeve Lowery, Antonio Omuro, Michael A. Postow, Dana Rathkopf, Alexander N. Shoushtari, Neerav Shukla, Martin H. Voss, Ederlinda Paraiso, Ahmet Zehir, Michael F. Berger, Barry S. Taylor, Leonard B. Saltz, Gregory J. Riely, Marc Ladanyi, David M. Hyman, Jos Baselga, Paul Sabbatini, David B. Solit, and Nikolaus Schultz. OncoKB: A Precision Oncology Knowledge Base. *JCO Precision Oncology*, 1(1):1–16, May 2017. ISSN 2473-4284. https://doi.org/10.1200/PO.17.00011. URL http://ascopubs.org/doi/full/10.1200/PO.17.00011.

Mostafa Karimi and Yang Shen. iCFN: an efficient exact algorithm for multistate protein design. *Bioinformatics*, 34(17):i811–i820, 2018.

Vikas Pejaver, Jorge Urresti, Jose Lugo-Martinez, Kymberleigh A. Pagel, Guan Ning Lin, Hyun-Jun Nam, Matthew Mort, David N. Cooper, Jonathan Sebat, Lilia M. Iakoucheva, Sean D. Mooney, and Predrag Radivojac. MutPred2: inferring the molecular and phenotypic impact of amino acid variants. *bioRxiv*, page 134981, May 2017. https://doi.org/10.1101/134981. URL https://www.biorxiv.org/content/early/2017/05/09/134981.

Damian Szklarczyk, John H. Morris, Helen Cook, Michael Kuhn, Stefan Wyder, Milan Simonovic, Alberto Santos, Nadezhda T. Doncheva, Alexander Roth, Peer Bork, Lars J. Jensen, and Christian von Mering. The STRING database in 2017: quality-controlled protein-protein association networks, made broadly accessible. *Nucleic Acids Research*, 45(D1):D362–D368, 2017. ISSN 1362-4962. https://doi.org/10.1093/nar/gkw937.
